## supplementary information for "A simple model for Behavioral Time Scale Synaptic Plasticity provides content addressable memory with binary synapses and one-shot learning"

November 24, 2024

### This PDF file includes

[1]Department of Computing, The Hong Kong Polytechnic University, Hong Kong SAR.

### S1. Equivalence of the BTSP rule from Eq. 1 with the rule proposed in Eq. 1 of [4] for the case of random plateau potentials

The BTSP rule from Eq. 1 of [4], which can be found on p.7 of the Supplement of that paper, is

$$\Delta w(t) = (1 - w(t))q_{\text{pre}}(t)q_{\text{post}}(t) - w(t)(q_{\text{pre}}(t)q_{\text{post}}(t-1) \vee q_{\text{pre}}(t-1)q_{\text{post}}(t)). \tag{S1}$$

Here, $w(t)$ represents a binary synaptic weight at time $t$, which can be either 0 or 1. $q_{\text{pre}}(t)$ denotes the binary indicator of spike signals or plateaus signals of presynaptic neurons, and $q_{\text{post}}(t)$ denotes the binary indicator of plateau signals of postsynaptic neurons. Their term $q_{\text{pre}}(t)$ corresponds to $x(t)$ and $q_{\text{post}}(t)$ corresponds to a binary indicator of a plateau potential in our notation.

We assume in this paper that plateau potentials occur at random time points in the postsynaptic neuron, as suggested by [2, 4]. This implies that if the presynaptic spike falls into the time window from -6s to +4s relative to the onset of a plateau potential, it falls according to the data of [5] and panel C of Fig. 1 with probability 0.5 into the LTP part of BTSP, and with probability 0.5 into the LTD part of BTSP (because both cover 5s of the relevant 10s time window). Hence, with probability 0.5, the first line of our rule is applied because the presynaptic spikes fall into the inner time window of Fig. 1 C (causing LTP if the current weight value is 0). This is equivalent to having $q_{pre}(t)q_{post}(t) = 1$ in the terminology of [4], which describes in their rule the case where LTP occurs if the current value is 0.

Dually, with probability 0.5, the second line of our BTSP rule (Eq.1) is applied because the presynaptic spike falls into the outer window of Fig. 1C with regard to the onset of the plateau potential in the postsynaptic neuron, causing LTD if the current weight value is 1. This is equivalent to having $q_{pre}(t)q_{pst}(t-1) = 1$ or $q_{pre}(t-1)q_{post}(t) = 1$, which describes in the rule of [4] the case where LTD is applied if the current weight value is 1. Hence, both rules are essentially equivalent. A minor difference arises in the interpretation of $q_{pre}(t)$ because the experimental data of [5] for BTSP in area CA1, that our rule models, do not consider –in contrast to [4]– the possibility of plateau potentials in the presynaptic neuron. Hence, $q_{pre}(t)$ should be interpreted in our context as only representing spikes of the presynaptic neuron.

Furthermore, there are small differences in the lengths of time windows for BTSP in [4], which models data from area CA3, and the lengths of time windows for BTSP in area CA1 of [5] that we model. But apart from these minor details, our BTSP rule in Eq. 1 is for the case of random plateau potentials at the given time windows mathematically equivalent to the BTSP rule of [4].

### S2. Experimental data on the emergence of place fields through BTSP from [1] can be reproduced with our BTSP rule

It was shown in Fig. 1A of [1] that BTSP supports the sudden emergence of a new place field while a rodent runs over a 180cm long repeating belt of a treadmill that was enriched with visual and tactile cues. These experimental data highlight the impact of BTSP on the time scale of seconds since synaptic weights from CA3 neurons that fire several seconds before or after the onset of a plateau potential in the CA1 neuron are strengthened; see also the symbolic plot in their Fig. 1F. We show in Fig.1D and F that our BTSP rule is able to reproduce these results, thereby demonstrating that it can model the experimentally observed impact of BTSP on the time scale of seconds.

We simulated repeated running of the mouse over a 180 cm long track at a slow running speed of 2 cm/s and at a fast running speed of 12cm/s. After completing one lap, the mouse is teleported back to the beginning of the track and carries out the next lap. Our model consists of 400 input neurons (modelling pyramidal cells in area CA3) and one output neuron (modelling a pyramidal cell in area CA1 to which the input neurons are synaptically connected). Initial weights have a value 1 with a probability 0.1. The input neurons have spatially tuned firing fields uniformly spaced across the linear track. As in Eq. 6 of the model of [5], we use the Gaussian function to model the firing rate of an input neuron with the maximum firing rate $R_{max} = 4$Hz. At the 10th lap, we randomly select a location and induce a plateau potential in the output neuron. We apply our stochastic BTSP rule to the synaptic weights from the input neurons to the output neuron for a total time of 10 seconds around the onset of the plateau potential.

For each input neuron that fires within this 10-second window, we modify the weight of its synapse to the output neuron once according to our BTSP rule (Eq. 1). Specifically, to apply our rule, we set the value of $x_i$ to 1 whenever the corresponding input neuron fires at least one spike within the 10-second window around the plateau potential. We apply the LTP part of the BTSP rule if the current weight is 0, otherwise the LTD part. Spikes of input neurons are generated through Bernoulli sampling, using a simulation time step of 0.5 second.

#### S3. Detailed estimates of properties of memory traces for overlapping memory items

This section provides detailed estimates for the memory traces of overlapping memory items. With the estimates of the probability for active synaptic weights (see Eqs. 22 and 23), we start by estimating the average firing probability of any output neuron after learning M overlapping patterns. In order to model the effect of common bits, we introduce the following two variables: (1) $c_w$, denoting the number of active connections from input neurons receiving common 1's to a given memory neuron after learning, and (2) $\hat{s}_j$, representing the number of weight updates for a synapse connecting neuron receiving common 1's during the learning. Recall the dependence of the active state of the weights on the parity of the number of weight updates: if $\hat{s}_j$ is odd, then the synapse will be in the active state after the learning, and thereby, the partial weighted sum of inputs from the common 1's is equal to $c_w$; otherwise, the synapse will be in the inactive state, contributing zero to the weighted sum.

By doing so, the introduction of $c_w$ and $\hat{s}_j$ can allow for a nuanced analysis of the firing probabilities, factoring in the distinct impact of common 1's on synaptic behaviour. We now evaluate the conditional firing probability of memory neurons to a given memory item post-learning as follows,

- For any $j_1 \in Q(\boldsymbol{x})$, we have

$$
\begin{aligned}
&\hat{p}_{f1}(\boldsymbol{x}, I, c_w, c) \\
&= P\big(z_{j_1}(\boldsymbol{x}) = 1 | j_1 \in Q(\boldsymbol{x}), \imath(\boldsymbol{x}) = I\big) \\
&= \sum_{l=0}^{M} P\big(z_{j_1}(\boldsymbol{x}) = 1, s_{j_1} = l | j_1 \in Q(\boldsymbol{x}), \imath(\boldsymbol{x}) = I\big) \quad \text{(by the total probability rule)} \\
&= \sum_{l=0}^{M} P(s_{j_1} = l) P\big(z_{j_1}(\boldsymbol{x}) = 1 | j_1 \in Q(\boldsymbol{x}), \imath(\boldsymbol{x}) = I, s_{j_1} = l\big) \quad \text{(by considering the parity of } \hat{s}_{j_1}) \\
&= \sum_{l=0}^{M} P(s_{j_1} = l) \Big( P\big(z_{j_1}(\boldsymbol{x}) = 1, \hat{s}_{j_1} \text{is even} | j_1 \in Q(\boldsymbol{x}), \imath(\boldsymbol{x}) = I, s_{j_1} = l\big) + P\big(z_{j_1}(\boldsymbol{x}) = 1, \hat{s}_{j_1} \text{is odd} | j_1 \in Q(\boldsymbol{x}), \imath(\boldsymbol{x}) = I, s_{j_1} = l\big) \Big) \\
&= \sum_{l=0}^{M} P(s_{j_1} = l) P(\hat{s}_{j_1} \text{is odd}) P\big(z_{j_1}(\boldsymbol{x}) = 1 | j_1 \in Q(\boldsymbol{x}), \imath(\boldsymbol{x}) = I, s_{j_1} = l, \hat{s}_{j_1} \text{is odd}\big) \\
&+ \sum_{l=0}^{M} P(s_{j_1} = l) P(\hat{s}_{j_1} \text{is even}) P\big(z_{j_1}(\boldsymbol{x}) = 1 | j_1 \in Q(\boldsymbol{x}), \imath(\boldsymbol{x}) = I, s_{j_1} = l, \hat{s}_{j_1} \text{is even}\big) \\
&= \sum_{l=0}^{M} P(s_{j_1} = l) \Big( \frac{1}{2}\big(1 - F(v_{th} - c_w, I - c, f_w \hat{p}_e(l))\big) + \frac{1}{2}\big(1 - F(v_{th}, I - c, f_w \hat{p}_e(l))\big) \Big) \\
&= \sum_{l=0}^{M} C_M^l f_q^l (1 - f_q)^{M-l} \Big( \frac{1}{2}\big(1 - F(v_{th} - c_w, I - c, f_w \hat{p}_e(l))\big) + \frac{1}{2}\big(1 - F(v_{th}, I - c, f_w \hat{p}_e(l))\big) \Big) \\
&\approx \sum_{l=0}^{M} \frac{(M f_q)^l}{2 e^{M f_q} l!} \Big( 2 - F\big(v_{th} - c_w, I - c, f_w \hat{p}_e(l)\big) - F\big(v_{th}, I - c, f_w \hat{p}_e(l)\big) \Big).
\end{aligned} \tag{S2}
$$

The last equation uses the Poisson approximation to the binomial probability for simplifying analysis.

- For any $j_2 \notin Q(\boldsymbol{x})$, we can obtain the estimate in a similar manner

$$
\begin{aligned}
&\hat{p}_{f2}(\boldsymbol{x}, I, c_w, c) \\
&= P(z_{j_2}(\boldsymbol{x}) = 1 | j_2 \notin Q(\boldsymbol{x}), \imath(\boldsymbol{x}) = I) \\
&= \sum_{l=0}^{M} P\big(z_{j_2}(\boldsymbol{x}) = 1, s_{j_2} = l, j_2 \notin Q(\boldsymbol{x}), \imath(\boldsymbol{x}) = I\big) \\
&= \sum_{l=0}^{M} P(s_{j_2} = l) P\big(z_{j_2}(\boldsymbol{x}) = 1 | j_2 \notin Q(\boldsymbol{x}), \imath(\boldsymbol{x}) = I, s_{j_2} = l\big) \quad \text{(by considering the parity of } \hat{s}_{j_2}) \\
&= \sum_{l=0}^{M} P(s_{j_2} = l) \Big( P\big(z_{j_2}(\boldsymbol{x}) = 1, \hat{s}_{j_2} \text{is even} | j_2 \notin Q(\boldsymbol{x}), \imath(\boldsymbol{x}) = I, s_{j_2} = l\big) + P\big(z_{j_2}(\boldsymbol{x}) = 1, \hat{s}_{j_2} \text{is odd} | j_2 \notin Q(\boldsymbol{x}), \imath(\boldsymbol{x}) = I, s_{j_2} = l\big) \Big) \\
&= \sum_{l=0}^{M} C_M^l f_q^l (1 - f_q)^{M-l} \Big( \frac{1}{2}\big(1 - F(v_{th}, I - c, f_w \hat{p}_o(l))\big) + \frac{1}{2}\big(1 - F(v_{th} - c_w, I - c, f_w \hat{p}_o(l))\big) \Big) \\
&\approx \sum_{l=0}^{M} \frac{(M f_q)^l}{2 e^{M f_q} l!} \Big( 2 - F\big(v_{th}, I - c, f_w \hat{p}_o(l)\big) - F\big(v_{th} - c_w, I - c, f_w \hat{p}_o(l)\big) \Big).
\end{aligned} \tag{S3}
$$

These estimations account for the firing probabilities of memory neurons post-learning. Combining Eqs.S2 and S3 gives the estimate of the average firing probability of memory neurons

$$\begin{aligned}
&\hat{p}_{f_{avg}}(\boldsymbol{x},c)\\
&=\sum_{I'=0}^{m-c}\sum_{c_w=0}^{c}P\big(\iota(\boldsymbol{x})=I'+c,c_w\big)P\big(u_j>v_{th}|\iota(\boldsymbol{x})=I'+c,c_w\big)\\
&=\sum_{I'=0}^{m-c}\sum_{c_w=0}^{c}P(\iota(\boldsymbol{x})=I'+c)P(c_w)\Big(f_q\hat{p}_{f1}\big(\boldsymbol{x},I'+c,c_w,c\big)+(1-f_q)\hat{p}_{f2}(\boldsymbol{x},I'+c,c_w,c)\Big)\\
&=\sum_{I'=0}^{m-c}\sum_{c_w=0}^{c}C_{m-c}^{I'}C_c^{c_w}\hat{f}_p^{I'}(1-\hat{f}_p)^{m-c-I'}f_w^{c_w}(1-f_w)^{c-c_w}\big(f_q\hat{p}_{f1}(\boldsymbol{x},I'+c,c_w,c)+(1-f_q)\hat{p}_{f2}(\boldsymbol{x},I'+c,c_w,c)\big),
\end{aligned}\tag{S4}$$

**Estimates of expected HD between memory traces for the original and masked memory items.** We use the firing probability of memory neurons to estimate the expected HD between memory traces for the original and masked memory items.

Unlike the estimate of randomly drawn memory items, the interaction of common bits and masking bits complicates the estimates. For simplification, we use the average number of masking common 1's, $cf_d$, to approximate the effects of the masking fraction on common 1's. On this basis, we evaluate $P(z_j(\boldsymbol{x})=1,z_j(\boldsymbol{x'})=0|\iota(\boldsymbol{x})=I)$ in two different cases, $j_1\in Q(\boldsymbol{x})$ and $j_2\notin Q(\boldsymbol{x})$.

- For any $j_1\in Q(\boldsymbol{x})$, we follow the derivation of Eq.S2 and have

$$\begin{aligned}
&\hat{p}_{c1}(j_1,\boldsymbol{x},\boldsymbol{x'},I,c_w)\\
&=P\big(z_{j_1}(\boldsymbol{x})=1,z_{j_1}(\boldsymbol{x'})=0|j_1\in Q(\boldsymbol{x}),\iota(\boldsymbol{x})=I\big)\\
&=\sum_{l=0}^{M}P\big(z_{j_1}(\boldsymbol{x})=1,z_{j_1}(\boldsymbol{x'})=0,s_{j_1}=l|j_1\in Q(\boldsymbol{x}),\iota(\boldsymbol{x})=I\big)\\
&=\sum_{l=0}^{M}P(s_{j_1}=l)P\big(z_{j_1}(\boldsymbol{x})=1,z_{j_1}(\boldsymbol{x'})=0|j_1\in Q(\boldsymbol{x}),\iota(\boldsymbol{x})=I,s_{j_1}=l\big)\\
&=\sum_{l=0}^{M}P(s_{j_1}=l)P(\hat{s}_{j_1}\text{is even})P\big(z_{j_1}(\boldsymbol{x})=1,z_{j_1}(\boldsymbol{x'})=0|j_1\in Q(\boldsymbol{x}),\iota(\boldsymbol{x})=I,s_{j_1}=l,\hat{s}_{j_1}\text{is even}\big)\\
&+\sum_{l=0}^{M}P(s_{j_1}=l)P(\hat{s}_{j_1}\text{is odd})P\big(z_{j_1}(\boldsymbol{x})=1,z_{j_1}(\boldsymbol{x'})=0|j_1\in Q(\boldsymbol{x}),\iota(\boldsymbol{x})=I,s_{j_1}=l,\hat{s}_{j_1}\text{is odd}\big)\\
&\approx\sum_{l=0}^{M}\frac{(Mf_q)^l}{2e^{Mf_q}l!}\Big(\big(1-F(v_{th}-c_w,I-c,f_w\hat{p}_e(l))\big)F\big(v_{th}-c_w(1-f_d),(I-c)(1-f_d),f_w\hat{p}_e(l)\big)\\
&+\big(1-F(v_{th},I-c,f_w\hat{p}_e(l))\big)F\big(v_{th},(I-c)(1-f_d),f_w\hat{p}_e(l)\big)\Big),
\end{aligned}\tag{S5}$$

where the handling of parity of $s_{j_1}$ follows the same as Eq.S2 and thus we simplify the derivation here.

- For any $j_2 \notin Q(\boldsymbol{x})$, we can derive the estimate in a similar manner:

$$\begin{aligned}
&\hat{p}_{c2}(j_2,\boldsymbol{x},\boldsymbol{x'},I,c_w)\\
&= P\big(z_{j_2}(\boldsymbol{x}) = 1, z_{j_2}(\boldsymbol{x'}) = 0 | j_2 \notin Q(\boldsymbol{x}), \iota(\boldsymbol{x}) = I\big)\\
&= \sum_{l=0}^{M} P\big(z_{j_2}(\boldsymbol{x}) = 1, z_{j_2}(\boldsymbol{x'}) = 0, s_{j_2} = l | j_2 \notin Q(\boldsymbol{x}), \iota(\boldsymbol{x}) = I\big)\\
&= \sum_{l=0}^{M} P(s_{j_2} = l) P\big(z_{j_2}(\boldsymbol{x}) = 1, z_{j_2}(\boldsymbol{x'}) = 0 | j_2 \notin Q(\boldsymbol{x}), \iota(\boldsymbol{x}) = I, s_{j_2} = l\big)\\
&= \sum_{l=0}^{M} P(s_{j_2} = l) P(\hat{s}_{j_2} \text{is even}) P\big(z_{j_2}(\boldsymbol{x}) = 1, z_{j_2}(\boldsymbol{x'}) = 0 | j_2 \notin Q(\boldsymbol{x}), \iota(\boldsymbol{x}) = I, s_{j_2} = l, \hat{s}_{j_2} \text{is even}\big)\\
&+ \sum_{l=0}^{M} P(s_{j_2} = l) P(\hat{s}_{j_2} \text{is odd}) P\big(z_{j_2}(\boldsymbol{x}) = 1, z_{j_1}(\boldsymbol{x'}) = 0 | j_2 \notin Q(\boldsymbol{x}), \iota(\boldsymbol{x}) = I, s_{j_2} = l, \hat{s}_{j_2} \text{is odd}\big)\\
&\approx \sum_{l=0}^{M} \frac{(Mf_q)^l}{2e^{Mf_q} l!} \Big(\big(1 - F(v_{th}, I-c, f_w\hat{p}_o(l))\big) F\big(v_{th}, (I-c)(1-f_d), f_w\hat{p}_o(l)\big)\\
&+ \big(1 - F(v_{th} - c_w, I-c, f_w\hat{p}_o(l))\big) F\big(v_{th} - c_w(1-f_d), (I-c)(1-f_d), f_w\hat{p}_o(l)\big)\Big).
\end{aligned} \tag{S6}$$

As the above Eqs.S5 and S6 do not depend on the specific positions of $j_1$ and $j_2$, we omit the indices $j_1$ and $j_2$ in the following derivations.

Combining Eqs.S5 and S6 gives the estimate of $E\big(HD(\boldsymbol{z}(\boldsymbol{x}), \boldsymbol{z}(\boldsymbol{x'}))\big)$

$$\begin{aligned}
&E\big(HD(\boldsymbol{z}(\boldsymbol{x}), \boldsymbol{z}(\boldsymbol{x'}))\big)\\
&= n \sum_{I=c}^{m} Pr(\iota(\boldsymbol{x}) = I) P\big(z_j(\boldsymbol{x}) = 1, z_j(\boldsymbol{x'}) = 0, \forall j | \iota(\boldsymbol{x}) = I\big)\\
&= n \sum_{I=c}^{m} Pr(\iota(\boldsymbol{x}) = I) \Big(f_q P\big(z_{j_1}(\boldsymbol{x}) = 1, z_{j_1}(\boldsymbol{x'}) = 0 | \iota(\boldsymbol{x}) = I, j_1 \in Q(\boldsymbol{x})\big)\\
&+ (1-f_q)\big(P(z_{j_2}(\boldsymbol{x}) = 1, z_{j_2}(\boldsymbol{x'}) = 0 | \iota(\boldsymbol{x}) = I, j_2 \notin Q(\boldsymbol{x}))\big)\Big)\\
&\approx n \sum_{I'=0}^{m-c} \sum_{c_w=0}^{c} C_{m-c}^{I'} C_c^{c_w} \hat{f}_p^{I'} (1-\hat{f}_p)^{m-c-I'} f_w^{c_w} (1-f_w)^{c-c_w} \Big(f_q \hat{p}_{c1}\big(\boldsymbol{x},\boldsymbol{x'},I'+c,c_w\big) + (1-f_q)\hat{p}_{c2}\big(\boldsymbol{x},\boldsymbol{x'},I'+c,c_w\big)\Big).
\end{aligned} \tag{S7}$$

$E\big(HD(\boldsymbol{z}(\boldsymbol{x}), \boldsymbol{z}(\boldsymbol{x'}))\big)$ provides the estimate of dissimilarity between the memory traces for the original and masked memory items, which was used in Fig. 4B.

**Estimates of expected HD between memory traces for different overlapping memory items.** To estimate the expected HD of memory traces between overlapping memory items, we use $\boldsymbol{x}_a$ and $\boldsymbol{x}_b$ to represent two different overlapping memory items. By using the estimate of firing probability for overlapping memory items, we can directly obtain an approximate estimate for this HD:

$$\begin{aligned}
&E\big(HD(\boldsymbol{z}(\boldsymbol{x}_a), \boldsymbol{z}(\boldsymbol{x}_b))\big)\\
&= \sum_{j=1}^{n} P\big(z_j(\boldsymbol{x}_a) = 1, z_j(\boldsymbol{x}_b) = 0\big) + P\big(z_j(\boldsymbol{x}_a) = 0, z_j(\boldsymbol{x}_b) = 1\big)\\
&\approx \sum_{j=1}^{n} P(z_j(\boldsymbol{x}_a) = 1) P(z_j(\boldsymbol{x}_b) = 0) + P(z_j(\boldsymbol{x}_a) = 0) P(z_j(\boldsymbol{x}_b) = 1)\\
&= 2n\hat{p}_{f_{avg}}(\boldsymbol{x}_a, c)\big(1 - \hat{p}_{f_{avg}}(\boldsymbol{x}_a, c)\big).
\end{aligned} \tag{S8}$$

For simplicity, we disregard the correlation arising from common bits in the above estimation. This approximation is suitable for cases where the fraction of common 1's is small, and the equal sign can be taken when the fraction of common 1's is 0. A more precise but intricate estimate can be obtained by expanding the estimation along $S(j)$, as exemplified by Eqs.S2 and S3. The estimated HD in Eq.S8 provides an approximation of the memory traces for different original memory items. By combining this estimate with the estimate in Eq.S7, we can directly obtain the estimate of the ratio of HD used in Fig. 4B.

### S4. Comparison of the performance of BTSP from Eq. (1) and core BTSP

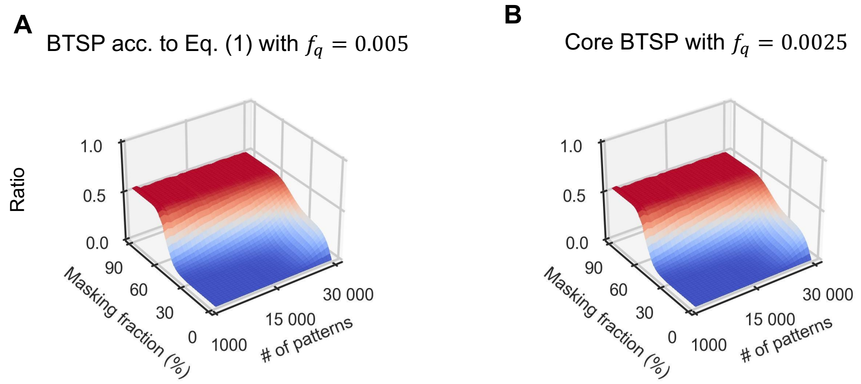

**Figure S1**: **Comparison of the performance of both versions of BTSP for recall with partially masked cues**. (**A**) Results for the BTSP version from Eq. (1), replotted from Fig. 3C. (**B**) Results of the same experiment with core BTSP, with $f_q = 0.0025$. Other parameter values remain the same. One sees that the performance of both versions of BTSP is virtually the same.

### S5. Distribution of inputs to memory neurons for BTSP and random projections

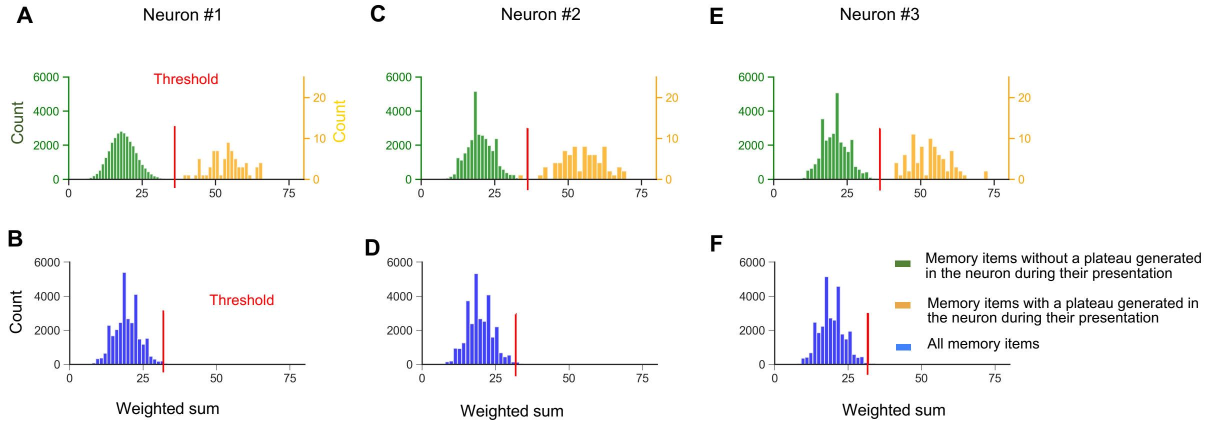

**Figure S2**: **Comparison of the distributions of inputs (=weighted sums) to 3 further samples of memory neurons for BTSP and RP; corresponding to Fig. 3 A, B for three further samples of memory neurons.** Panels (**A**,**C** and **E**) Same BTSP experiment as for Fig. 3 B, for 3 further randomly drawn memory neurons. Panels (**B**,**D** and **F**) Same RP experiment as for Fig. 3 A, for 3 further randomly drawn memory neurons.

#### 114 S6. Comparison of recall results for perturbed and partially masked cues

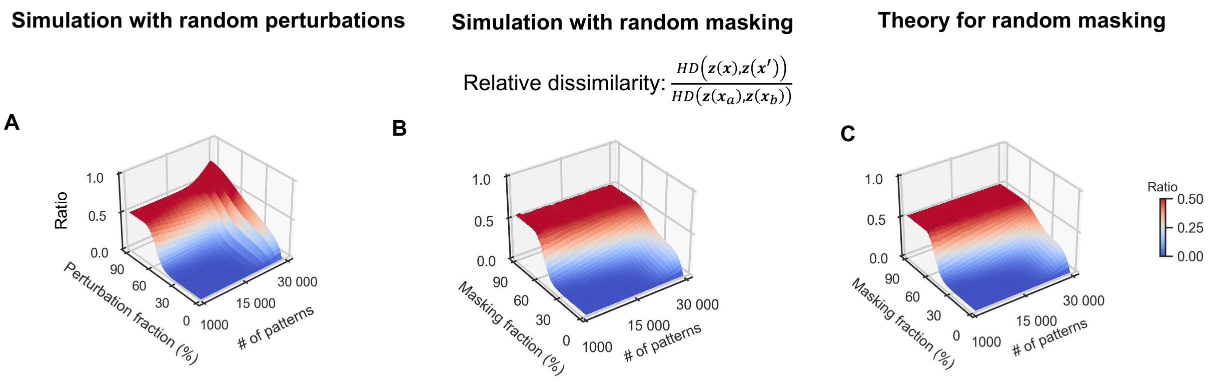

**Figure S3**: **Comparison of results for recall with perturbed and partially masked cues**. (**A**) Recall of memory traces with perturbed cues, where not only 1's were replaced by 0's, but also 0's were replaced by 1's; keeping, however, the average number of 1's the same as in the original input patterns. For any given perturbation fraction $f'_d$, it is assumed that each 1 in an input pattern is switched to 0 with a probability equal to $f'_d$, and each 0 is switched to 1 with a probability equal to $(mf'_d f_p)/(m(1-f_p)) = f'_d f_p/(1-f_p)$. The optimization process and parameter setups remain the same as in Fig. 3C, except that the thresholds were separately optimized for the two types of tests with perturbed and partially masked input patterns. The theoretical results for panel C are derived from Eqs. (17) and (21). One sees that the impact of perturbation on recall performance is virtually the same as for partially masked input patterns.

#### 115 S7. Impact of $f_q$ on recall performance of BTSP

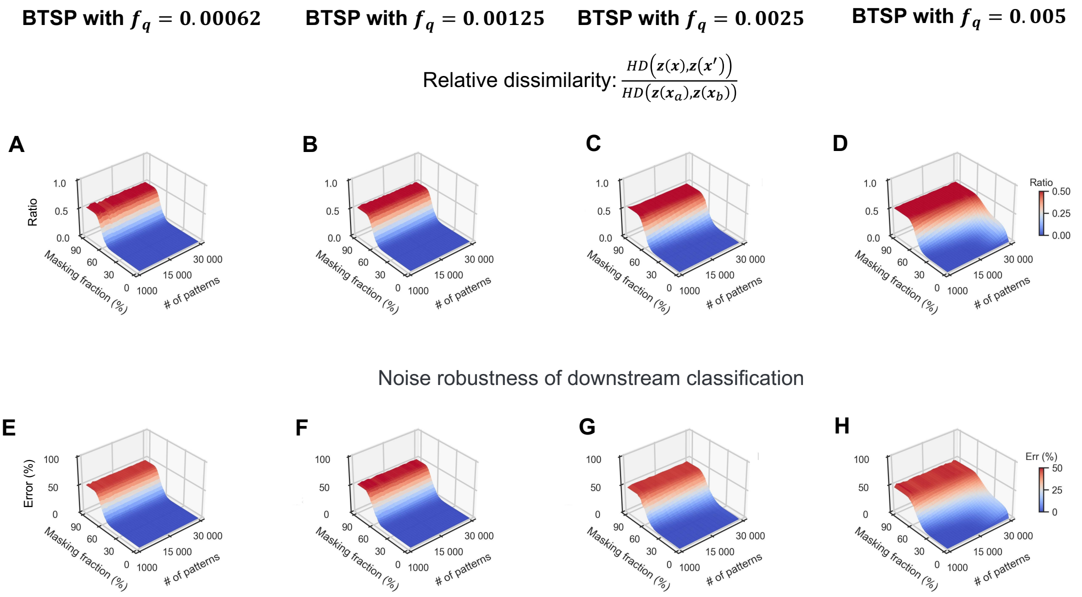

**Figure S4**: **Comparison of the memory traces created by BTSP with reduced values of $f_q$.** Panels **D**And **H** for the default value of $f_q$ are copied from Fig.3C and F. Panels (**A**) to (**C**) show results of the same experiments but with lower values of $f_q$. One sees that the performance improves a little bit further for lower values of $f_q$.

#### 116 S8. Impact of the probability of the LTD part of BTSP

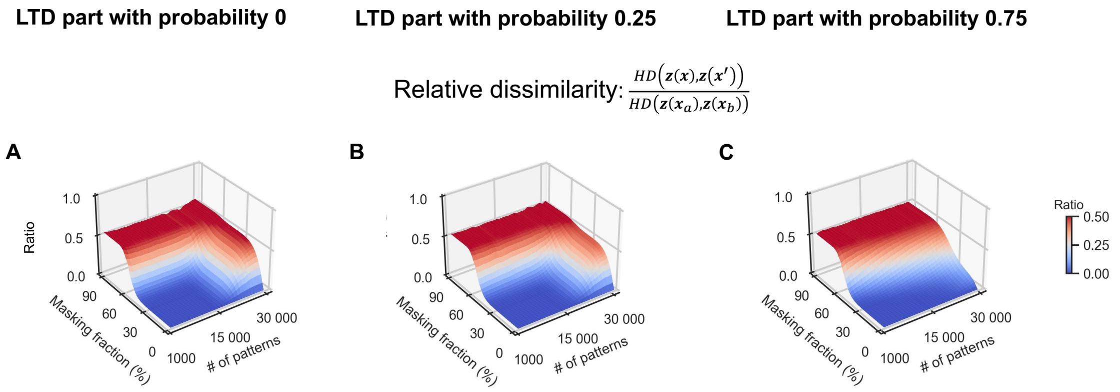

**Figure S5**: **Impact of the probability of the LTD part of BTSP on relative dissimilarity; compare with Fig. 3C** (**A**-**C**). We changed the probability of LTP and LTD to values between 0.25 and 0.75 and repeated the experiment of Fig.3C. Other parameter values remain the same as there. One sees that the performance increases somewhat for larger probabilities of LTD. Note that a larger probability of LTD also reduces the number of weights with values 1 in the memory system. While the experimental data of [5] suggest a probability around 0.5, a larger probability of LTD could model the impact of separate weight decay processes.

#### 117 S9. Dependence of CAM performance with BTSP on the value of $f_q$

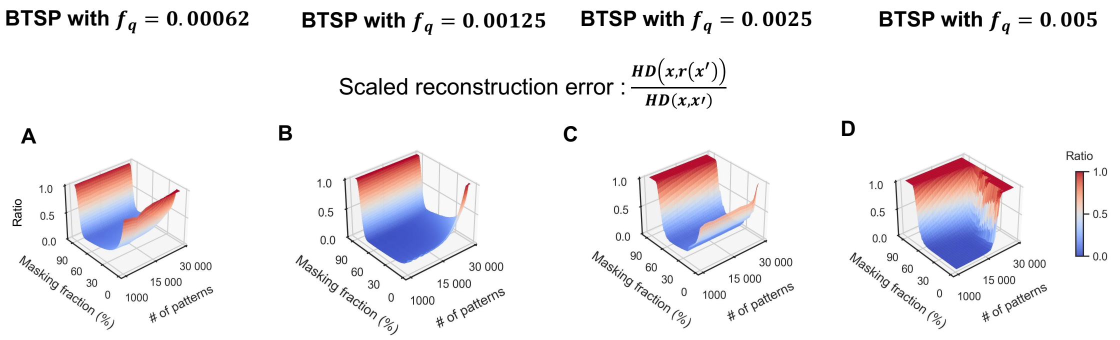

**Figure S6**: **Comparison of the input reconstruction capability of BTSP with smaller values of** $f_q$. Panel D shows again the performance for the default value of $f_q$, copied from Fig.5D. (**A**-**C**) Corresponding CAM performance results with lower values of $f_q$. One sees that the performance is improved with a lower value of $f_q$.

### S10. Scaling up the size of input patterns to the size of area CA3 in the human brain

We have so far examined memory structures for a fixed size of the model that we introduced at the beginning, which corresponds to about 1/100 of the number of neurons in corresponding structures of the human brain. We address here the question how the memory capacity of the BTSP model changes when we use it to store 100 times larger input patterns that roughly correspond to the size of area CA3 in the human brain. Simultaneously we multiply the probability $f_p$ that an input neuron is active for a memory item $\boldsymbol{x}$ with the factor 1/100 so that on average the same number of input neurons are active for a memory item $\boldsymbol{x}$. We keep the number $n$ of memory neurons the same as in our default setting.

A similar scaling up of the size of input features arises when one takes a more complex neuron model into account, even if one stays at the scale of the mouse brain. The recent review [3] suggests in Fig. 4 a qualitative rule for synaptic integration in a pyramidal cell with dendrites whereby each segment of its dendritic arbor should be viewed as an independent small integration zone that generates an NMDA spike if several synapses in this segment are co-active. If one assumes for simplicity that there are exactly 2 synapses in each dendritic segment, and that this segment generates an NMDA spike if and only if these two synapses are co-active, one arrives at a virtual input space that consists of all pairs of presynaptic neurons that form synapses at a common dendritic segment. This virtual input space is in general much larger. At the same time, the sparseness of activity is decreased by a corresponding factor because both synapses at a dendritic segment are more rarely co-active than individual presynaptic neurons.

We find that this change in the pattern size has a drastic impact on the memory capacity and recall capability with partial cues, see Fig. **S7**B, compared with Fig. 3C. Due to computational resource limitations, simulations were only conducted for up to 20,000 patterns. Since there is almost perfect agreement between theoretical predictions and simulation outcomes for up to 20,000 patterns, comparing Fig. **S7** A and B indicates that the theoretical predictions appear to be trustworthy also for larger numbers of input patterns. Results of these theoretical predictions for up to 800,000 input patterns are shown in Fig. **S7**B.

Fig. **S7**A and B show that in this setting with 100 times larger input patterns, the attractor properties of memory traces created through BTSP become even more pronounced: Good recall is possible even when up to 2/3 of the bits of a memory item have been masked in a recall cue. Hence, both the capacity and the robustness of recall are substantially improved in this setting, which is arguably more realistic from the biological perspective. This result also suggests that one can build with the help of BTSP powerful CAM models in neuromorphic hardware that can store up to 800,000 patterns, and support robust recall even when more than 2/3 of the features of a memory item are missing in a recall cue.

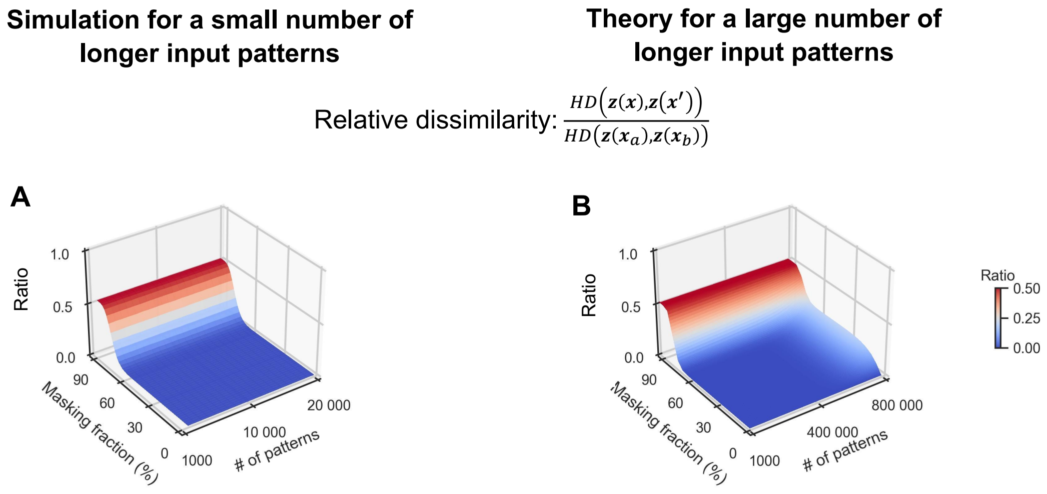

**Figure S7**: **Scaling-up the number and sizes of input patterns.** (**A**) Results of numerical experiments are shown for up to 20,000 input patterns that are each 100 times longer, but where the probability that an input bit has value 1 is 1/100 times smaller than the expected number of 1s in an input pattern remains the same. This scaling result enables successful recall even when up to two-thirds of the bits in a memory item are masked in the recall cue. (**B**) Theoretical predictions for the same type of input patterns derived from Eqs. 17 and 21, but for up to 800,000 input patterns. One sees excellent agreement with the simulation results of A for smaller pattern numbers. These theoretical results predict very robust recall with partial cues where up to 2/3 of bits are masked, for up to 800,000 patterns, i.e., for much larger numbers of input patterns.

### S11. Comparison of input completion capabilities of different memory models

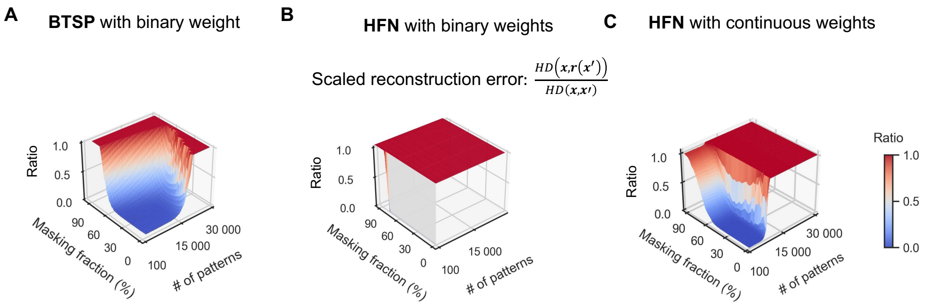

**Figure S8**: **Comparison of input completion capabilities of different memory models; compare with Fig.5D**. (**A**) A quantitative analysis of the input reconstruction capability of the BTSP model where the number *n* of neurons in the memory layer was reduced from 39,000 to 25,000, which is the number of neurons in the input layer and, therefore, also the bit length of a memory item. The performance is only affected very little by this reduction. We had chosen n > m in our default setting to reflect the relative sizes of areas CA3 and CA1 in the brain. We show here that the same memory capacity can be achieved when *n* is lowered to the value of *m*, i.e., to the length 25,000 of each memory item. These values may be seen as providing a fairer comparison to HFNs without hidden units. (**B**) shows a quantitative analysis of the input reconstruction capability of the HFN model with binary weights for the same ensemble of random input patterns.The binarization strategies for HFN models are elaborated in Figure **S9**.(**C**) shows a quantitative analysis of the input reconstruction capability of the HFN model with continuous weights.

### S12. CAM properties of HFNs with binarized weights

Since CAM systems with continuous weights are hard to emulate in neuromorphic hardware, the question arises whether it makes sense to use HFNs with binarized weights. However, a drastic drop in memory capacity after rounding of weights had been demonstrated in [6] for the case of densely coded memory items. Hence we wondered whether this drastic drop in memory capacity also occurs for the sparse input patterns and incomplete network connectivity on which we have focused.

We employed a fixed threshold to binarize the HFN's weights. We set the weight values to 0 or $1/m$, thereby aligning the range of binarized values with that of the original weight matrix, $\boldsymbol{W}^H$, defined in Eq.(31). Three different binarization threshold values were evaluated for $\boldsymbol{W}^H$: (1) 0, (2) the median of $\boldsymbol{W}^H$, and (3) the mean of $\boldsymbol{W}^H$. For each binarization threshold, we searched for the value of $v_{th}^H$ from $[0.0, 0.02]$ with the minimum step 0.001 with the same optimization criterion as used for the continuous weights (see *Methods*). Results for input completion with different binarization thresholds and different $v_{th}^H$ are provided in Fig. **S9**. The resulting optimal threshold, $v_{th}^H = 0.001$, was adopted in the main text.

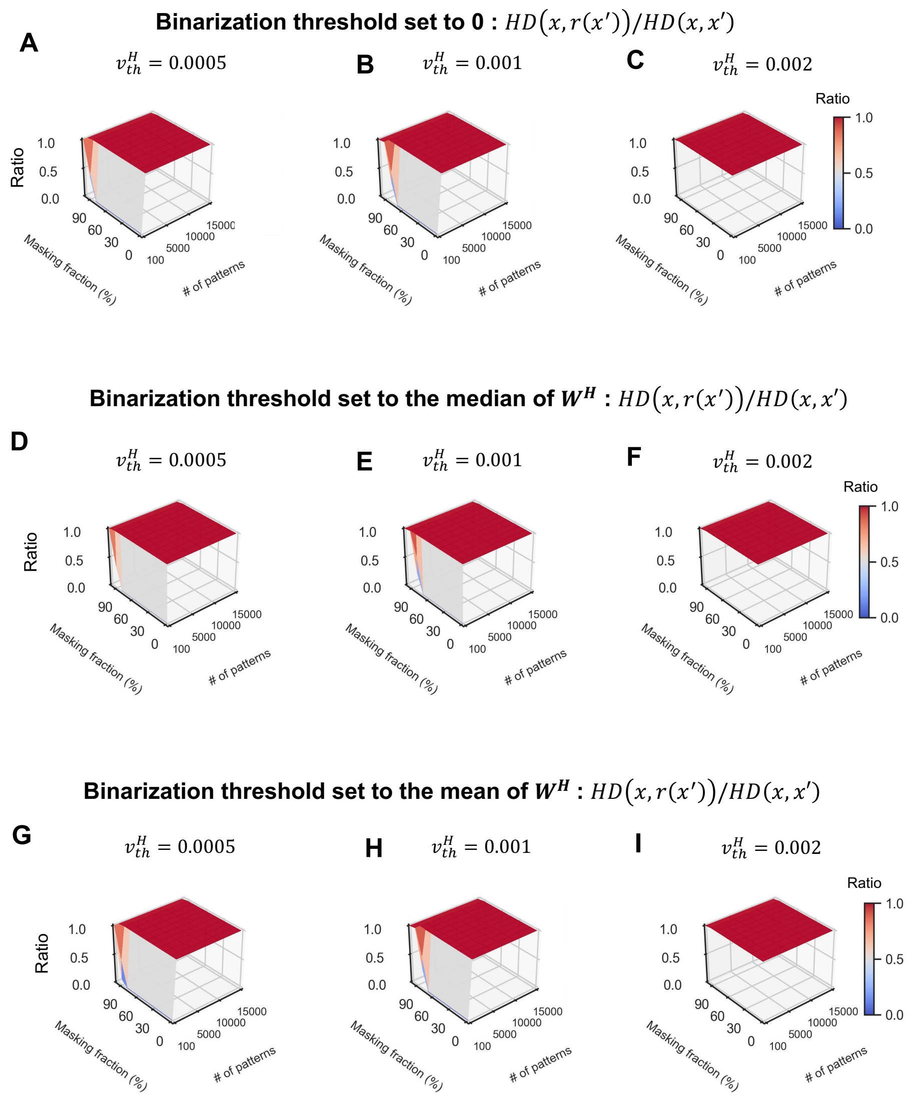

**Figure S9**: **Analysis of CAM performance of a HFN with various binarization strategies for synaptic weights and different values of thresholds $v_{th}^H$ of neurons; compared with Fig.5E**. (**A**-**C**) Simulation results of HFN using a binarization threshold of 0 with different $v_{th}^H$ values. (**D**-**F**) Simulation results of HFN using a binarization threshold equal to the median of the weight matrix $\boldsymbol{W}^H$ (defined in Eq.(31)) with various $v_{th}^H$ values. (**G**-**I**) Simulation results of HFN using a binarization threshold equal to the mean of the weight matrix $\boldsymbol{W}^H$ with different $v_{th}^H$ values. One sees that no decent CAM performance can be achieved by HFNs with binary weights for any of the parameters that we tried.

### Supplementary References

[1] Katie C Bittner, Aaron D Milstein, Christine Grienberger, Sandro Romani, and Jeffrey C Magee. Behavioral time scale synaptic plasticity underlies ca1 place fields. *Science*, 357(6355):1033–1036, 2017.

[2] Christine Grienberger and Jeffrey C Magee. Entorhinal cortex directs learning-related changes in ca1 representations. *Nature*, pages 1–9, 2022.

[3] Matthew E Larkum. Are dendrites conceptually useful? *Neuroscience*, 489:4–14, 2022.

[4] Yiding Li, John J Briguglio, Sandro Romani, and Jeffrey C Magee. Mechanisms of memory-supporting neuronal dynamics in hippocampal area ca3. *Cell*, 2024.

[5] Aaron D Milstein, Yiding Li, Katie C Bittner, Christine Grienberger, Ivan Soltesz, Jeffrey C Magee, and Sandro Romani. Bidirectional synaptic plasticity rapidly modifies hippocampal representations. *Elife*, 10:e73046, 2021.

[6] Mikhail S Tarkov. Hopfield associative memory with quantized weights. In *Advances in Neural Computation, Machine Learning, and Cognitive Research II: Selected Papers from the XX International Conference on Neuroinformatics, October 8-12, 2018, Moscow, Russia*, pages 91–97. Springer, 2019.
